# Perversely expressed long noncoding RNAs can alter host response and viral proliferation in SARS-CoV-2 infection

**DOI:** 10.1101/2020.06.29.177204

**Authors:** Rafeed Rahman Turjya, Md. Abdullah-Al-Kamran Khan, Abul Bashar Mir Md. Khademul Islam

## Abstract

**Background:** Since December 2019, the world is experiencing an unprecedented crisis due to a novel coronavirus, SARS-CoV-2. Owing to poor understanding of pathogenicity, the virus is eluding treatment and complicating recovery. Regulatory roles of long non-coding RNAs (lncRNAs) during viral infection and associated antagonism of host antiviral immune responses has become more evident in last decade. To elucidate possible functions of lncRNAs in the COVID-19 pathobiology, we have utilized RNA-seq dataset of SARS-CoV-2 infected lung epithelial cells.

**Results:** Our analyses uncover 21 differentially expressed lncRNAs whose functions are broadly involved in cell survival and regulation of gene expression. By network enrichment analysis we find that these lncRNAs can directly interact with differentially expressed protein-coding genes *ADAR, EDN1, KYNU, MALL, TLR2* and *YWHAG*; and also *AKAP8L, EXOSC5, GDF15, HECTD1, LARP4B, LARP7, MIPOL1, UPF1, MOV10* and *PRKAR2A*, host genes that interact with SARS-CoV-2 proteins. These genes are involved in cellular signaling, metabolism, immune response and RNA homeostasis. Since lncRNAs have been known to sponge microRNAs and protect expression of upregulated genes, we also identified 9 microRNAs that are induced in viral infections; however, some lncRNAs are able to block their usual suppressive effect on overexpressed genes and consequently contribute to host defense and cell survival.

**Conclusions:** Our investigation determines that deregulated lncRNAs in SARS-CoV-2 infection are involved in viral proliferation, cellular survival, and immune response, ultimately determining disease outcome and this information could drive the search for novel RNA therapeutics as a treatment option.

## Background

By mid-May 2020, the confirmed cases of COVID-19 have crossed 4 million (1), making it one of the largest pandemics in modern times. Since the first reported case at Wuhan, China in December 2019 (2), this viral disease has been responsible for more than 300,000 deaths worldwide (1), and the number is steadily rising. The causative agent behind the disease is a novel beta-coronavirus (3), which has been named SARS Coronavirus 2 i.e. SARS-CoV-2 due to its similarity to the earlier SARS-CoV first detected in 2002 (4). The rapid spread of the virus, the ever-increasing death toll, and absence of a sufficient treatment strategy has affected societies and economies all over the globe.

The molecular mechanism of SARS-CoV-2 is complex and interrelated with host mechanisms, as is common with most pathogenic viruses (5). It is possible that infected lung epithelial cells trigger innate immune pathways, leading to immune effector cells releasing high levels chemokines and proinflammatory cytokines and resultant unconfined or uncontrolled systemic inflammatory response leads to fatality (6, 7). Recent studies also indicate the virus may cause viral sepsis (8) and infection can lead to deregulation in blood coagulation (9, 10). But no one conjecture has been able to formulate a clear and concise explanation of how the virus spreads so effectively and affects so profoundly. With emergence of more and more data, new dimensions arise as possible modes of pathogenesis and progression of infection.

Non-coding RNAs (ncRNAs), RNAs that do not code for any proteins, are key players in the regulation of gene expression and influence the interplay involved in host defense mechanisms (11). Regulatory ncRNAs, such as microRNAs (miRNAs) and long non-coding RNAs (lncRNAs) act as important regulators of the cellular antiviral response. Consequently, viruses have been found to utilize cellular ncRNA to evade immune response and exploit cellular machinery to their advantage (12).

LncRNAs are a type of non-coding RNA having a size of more than 200 nts that can function as primary or spliced transcripts (13). LncRNAs have become increasingly crucial in explaining cellular processes and understanding molecular progression of diseases. They may regulate gene expression through epigenetic modification of chromatin structure, transcriptional control, regulation of gene transcription via direct binding or transcription factor recruitment and post-transcriptional processing through protein-RNA interaction (14–16). Additionally, lncRNAs may function as competing endogenous RNAs (17) to function as regulators of microRNA targeting of genes involved in important pathways.

Previous studies have found lncRNAs to be involved in viral infection and subsequent host response (18–20). A wide selection of lncRNAs get aberrantly modulated in many viral infections like-HSV, Influenza, HIV, HSV etc (21). Upon viral infections, dysregulation of cellular lncRNAs occur which in turn abnormally regulates several host processes resulting in the progression of the viral infection (20). Apart from the general host routes, abnormalities in the expression of lncRNAs mainly affect host’s different antiviral innate immune responses, particularly interferon signaling and IFN-stimulated genes (ISGs) (20, 21). Even in SARS-CoV infection, Peng et al. showed that differential expression of lncRNAs could aberrantly regulate several host responses along with the innate immune signaling (22), which suggests a similar deregulation pattern of lncRNAs could also occur in SARS-CoV-2 infection.

Blanco-Melo et al. (23) investigated the host response to SARS-CoV-2 by infecting primary human lung epithelium (NHBE) cells and A549 alveolar cell lines with the virus and performing RNA-seq analysis to identify differentially expressed (DE) genes. But no such study yet concluded the possible outcomes of the deregulated lncRNAs in SARS-CoV-2. In our present study we have identified lncRNAs that are differentially expressed in SARS-CoV-2 infected cell’s transcriptome compared to uninfected cells, and then correlated the putative effects of the deregulated lncRNAs in the tug-of-war between SARS-CoV-2 and the host. We have also investigated the possible aftermaths of lncRNA deregulation in COVID-19 disease pathobiology.

## Results

### lncRNAs differentially modulated in SARS-CoV-2 infection

To investigate the probable deregulation of lncRNAs in SARS-CoV-2 infection, firstly we have analyzed the 24 hours post-infection transcriptome data of SARS-CoV-2 infected NHBE cells. Analyzing the RNA-seq data led to identification of 687 DE genes and intriguingly, amongst those DE genes, we discovered 21 lncRNA genes, 9 of them upregulated while 12 were downregulated (Figure 1).

**Figure 1:**
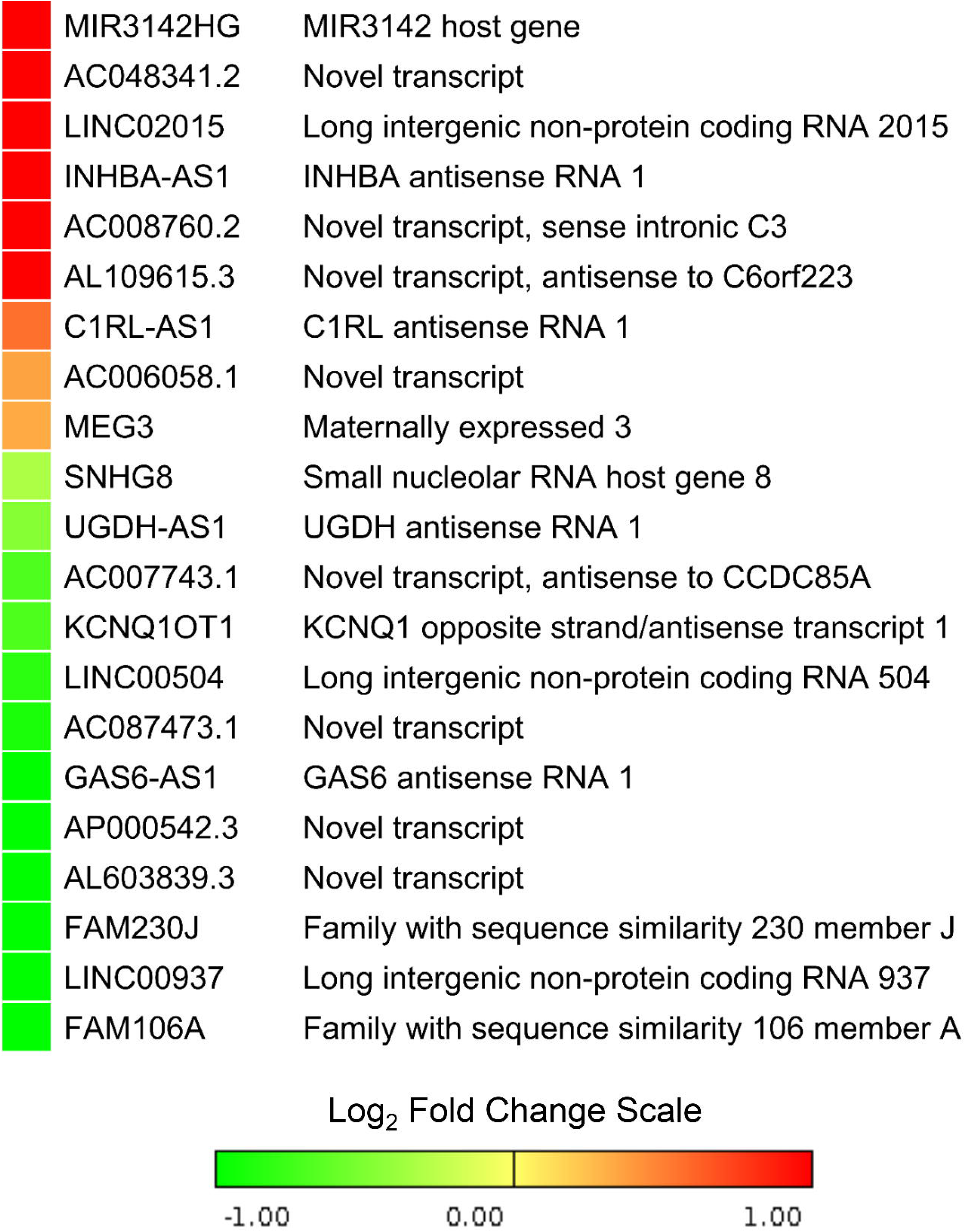
Differentially expressed lncRNAs in SARS-CoV-2 infection. Gene expressions are presented in Log2 fold change (compared to uninfected control cells) value color coded heatmap. Color towards red indicates more upregulation and color towards green indicates further downregulation, while yellow color indicates absence of differential expression.

### lncRNAs interact with several differentially modulated proteins in SARS-CoV-2 infected cells

We now sought to elucidate the putative effects of these deregulated lncRNAs in SARS-CoV-2 infection. In order to achieve that, we built a network of the deregulated lncRNAs along with their interacting protein coding target genes. Among the differentially expressed protein-coding genes, 6 were found to interact with the DE lncRNAs which (Table 1). There are direct RNA-RNA interactions for 4 genes, and RNA-protein interactions for the rest (Figure 2). These lncRNA interacting protein coding genes have potential roles during the viral infections (Table 1).

**Table 1:**
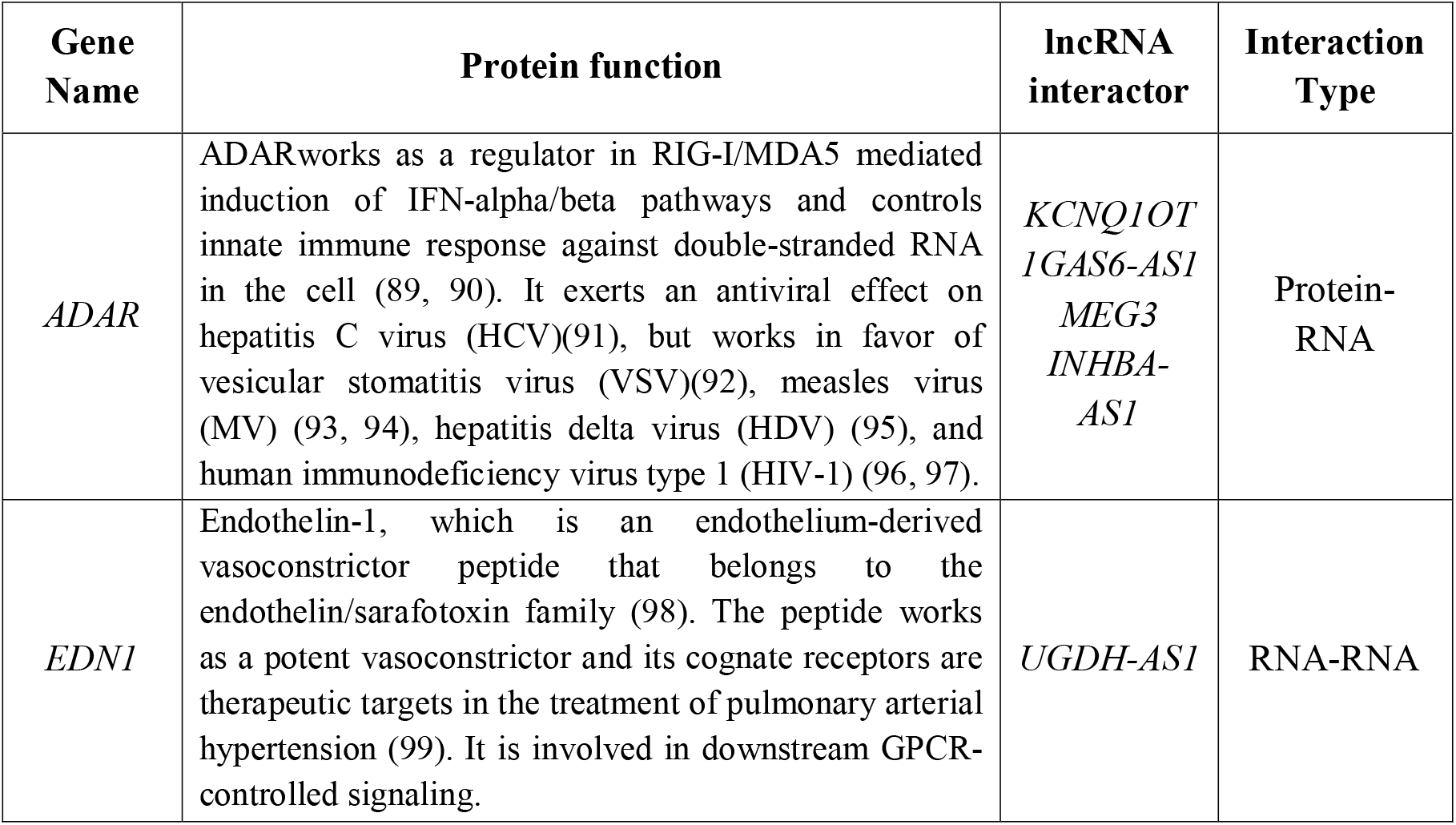
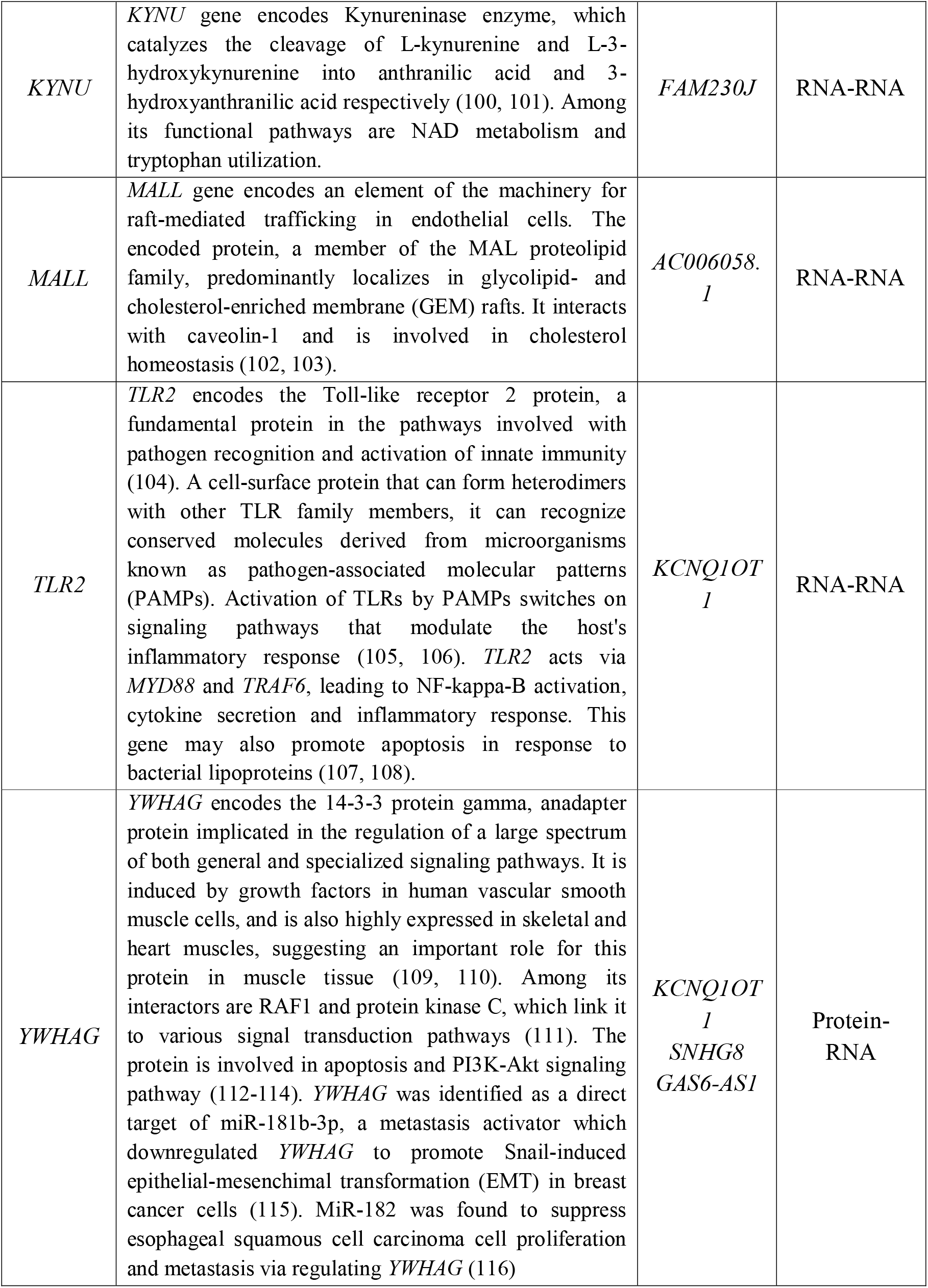
DE genes, their encoded protein’s functions and associated interacting lncRNAs.

**Figure 2:**
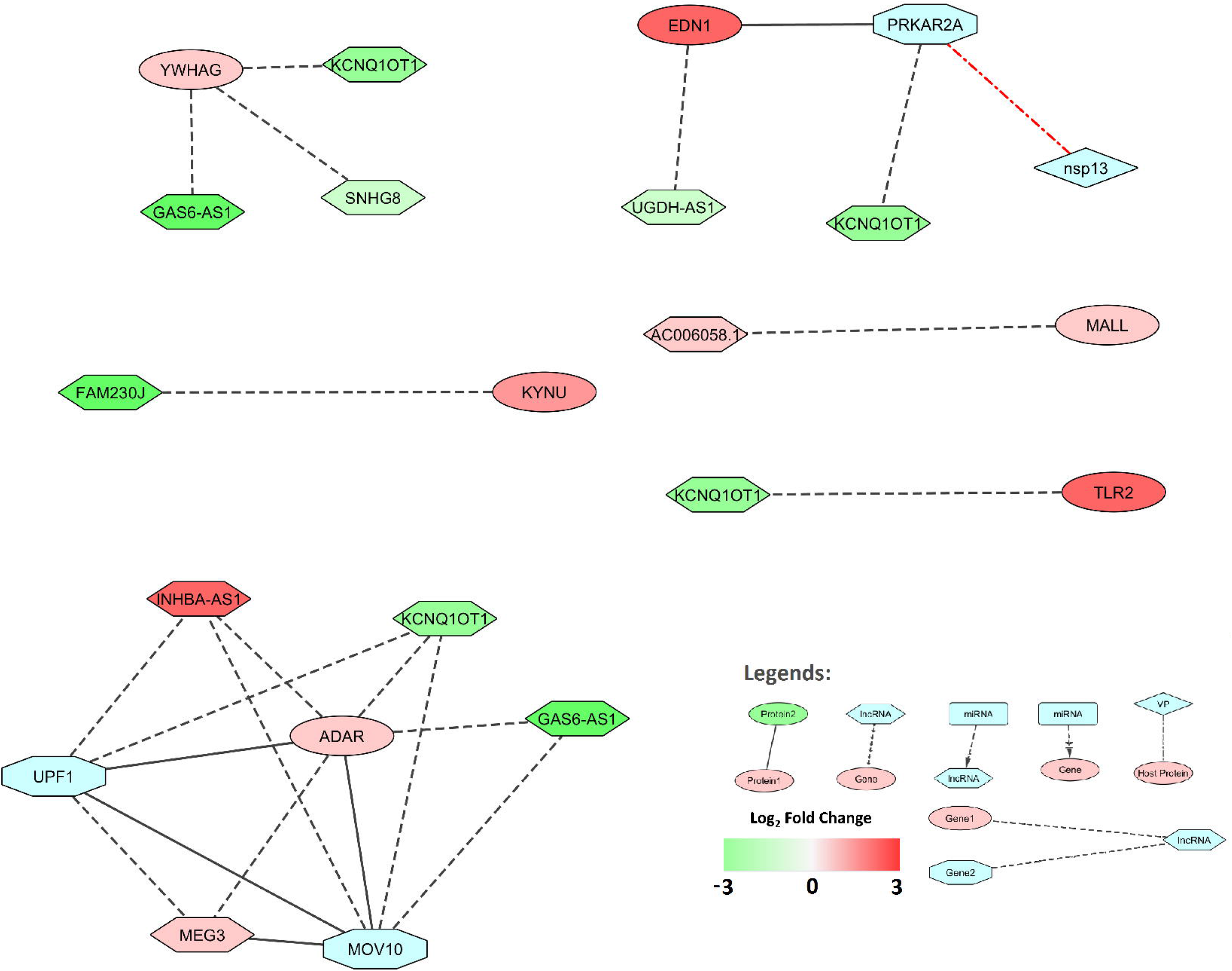
Differentially expressed lncRNAs that interact with differentially expressed protein-coding Genes. Node shape legends: ellipse: DE gene, hexagon: lncRNA, diamond: viral protein, Octagon: viral protein interactor, rectangle: microRNA; types of edges are- dot-dash: VP-host protein interaction; dash: lncRNA-gene interaction, dash-arrow: miRNA-gene interaction; solid: protein-protein interaction; sine wave: ceRNA; separate arrows: alternate target. Log2 Fold change color scale is as depicted in Figure 1.

Upregulated *ADAR* interacts directly with downregulated *KCNQ1OT1* and *GAS6-AS1*, and upregulated *MEG3* and *INHBA-AS1* (Figure 2). ADAR also interacts with UPF1 and MOV10 – two proteins which were also interactors of other DE lncRNAs. *EDN1* is found to be upregulated in SARS-CoV-2 infection, which interacts directly with downregulated *UGDH-AS1* (Figure 2). EDN1 can also interact with PRKAR2A, a protein that interacted with *KCNQ1OT1*. *KYNU* mRNA interacts directly with *FAM230J*, which was downregulated (24). *MALL* gene was found upregulated in SARS-CoV-2 infection. This mRNA interacts directly with *AC006058.1*, which was also upregulated. Upregulated *TLR2* interacts directly with downregulated lncRNA *KCNQ1OT1* (Figure 2). YWHAG protein interacts directly with *KCNQ1OT1*, *SNHG8*, *GAS6-AS1*, all of which were downregulated (Figure 2).

### Protein interactors of SARS-CoV-2 viral proteins also interacted with DE lncRNAs

Gordon et al. investigated possible protein-protein interactions (PPI) between SARS-CoV-2 proteins and host proteins and identified 332 high-confidence interactions (25). We wanted to stratify whether these interacting proteins can be also interact with the DE lncRNAs in SARS-CoV-2 infection. We have constructed a network with the DE lncRNAs and their interacting host proteins that also bind with viral proteins. Among these 332 proteins, we found 10 such proteins to interact with the DE lncRNAs (Table 2). Among them, 7 proteins interacted at protein-RNA level, whereas the rest were RNA-RNA interactions. SARS-CoV-2 proteins M (Membrane), N (Nucleocapsid), Nsp8, Nsp12, Nsp13, and ORF8 were found to interact with these host proteins (Figure 3).

**Table 2:**
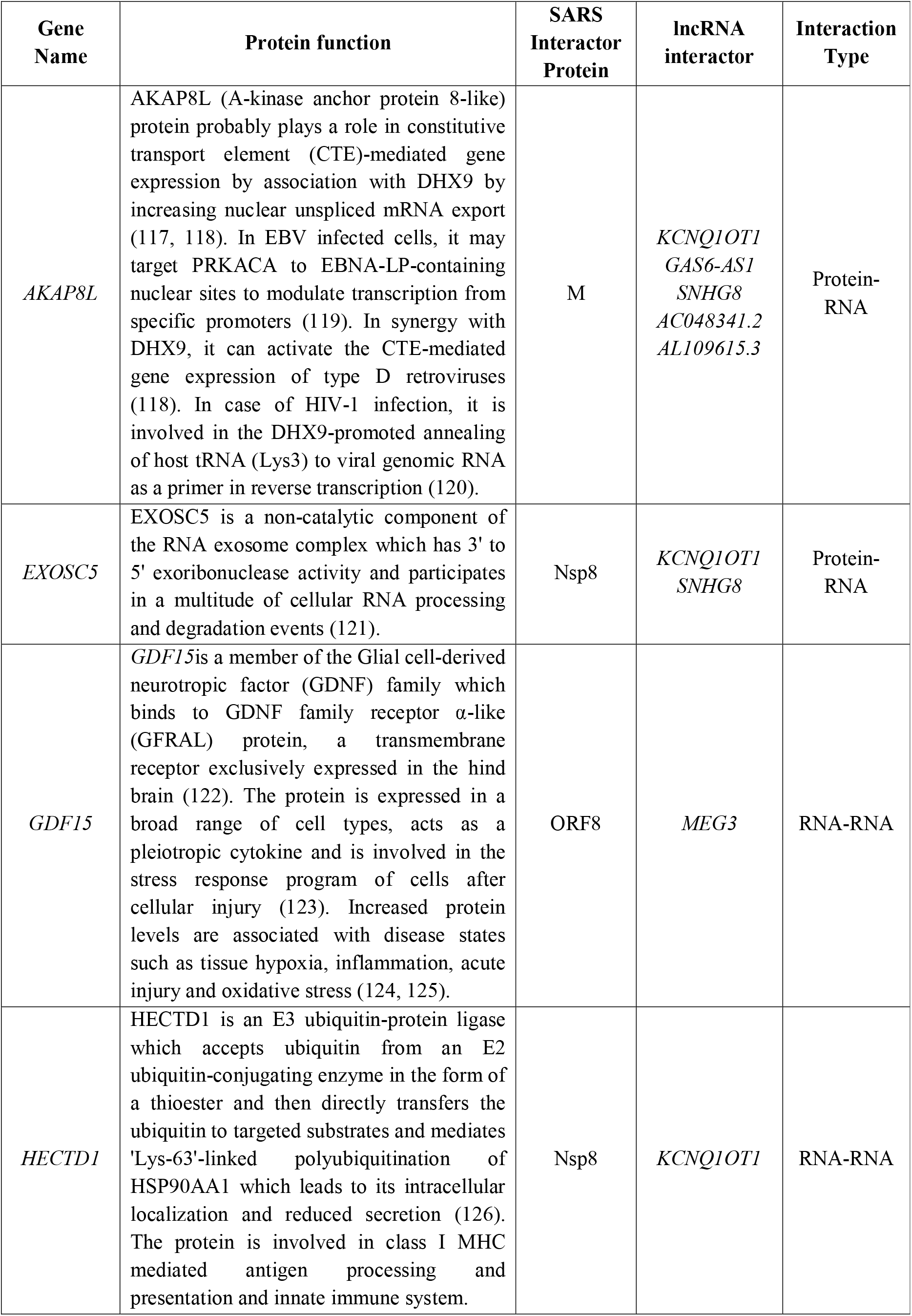
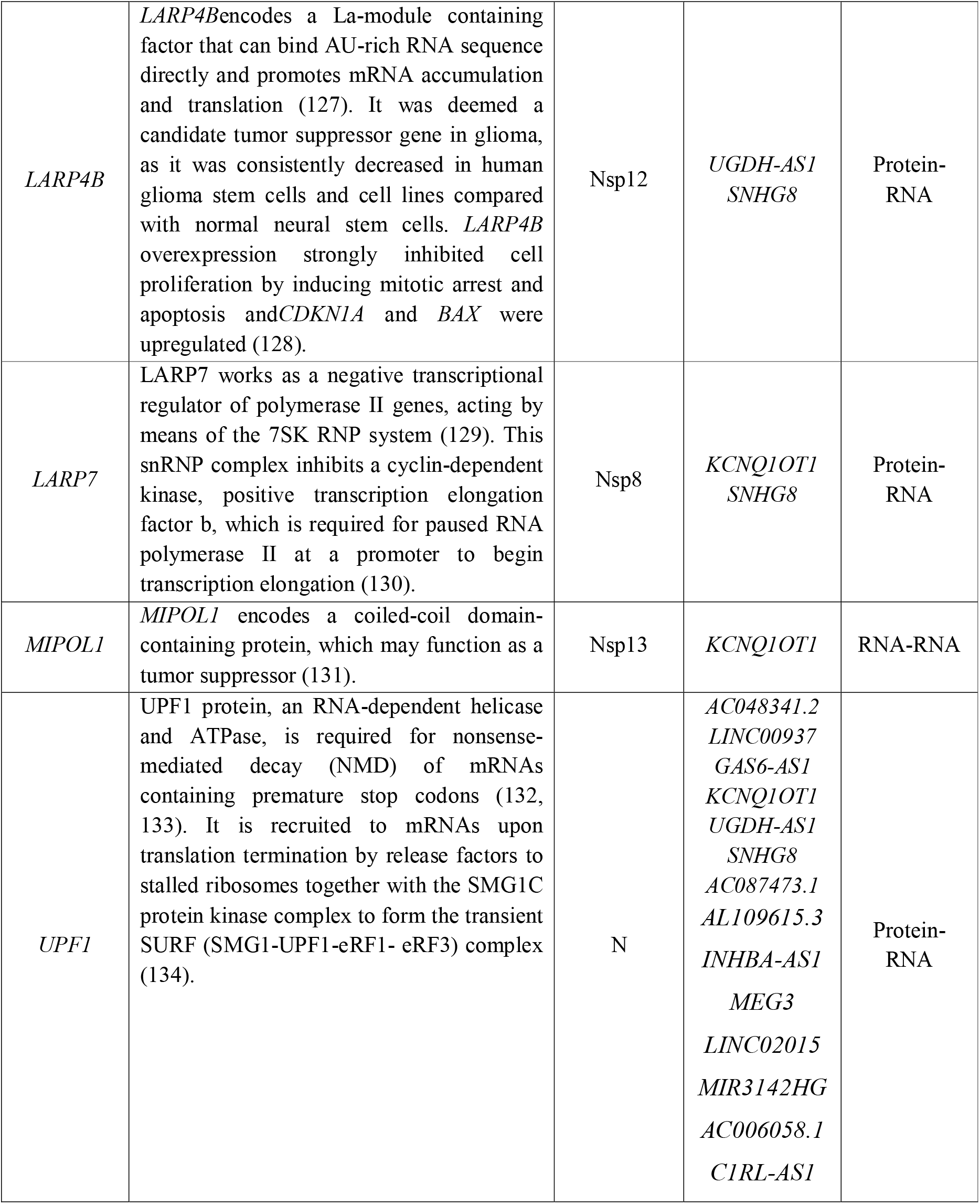
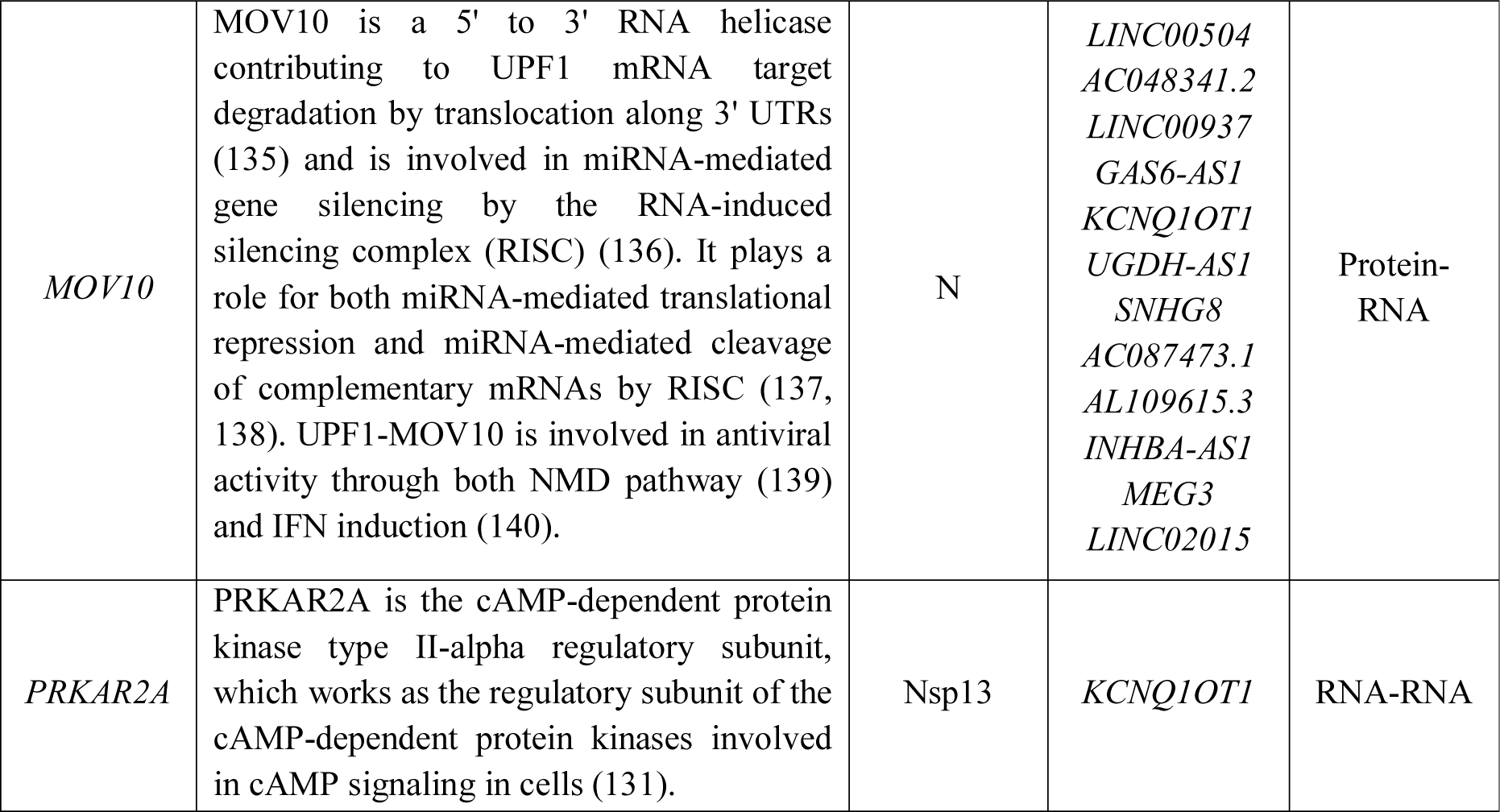
SARS-CoV-2 interactor host proteins and associated interacting lncRNAs.

**Figure 3:**
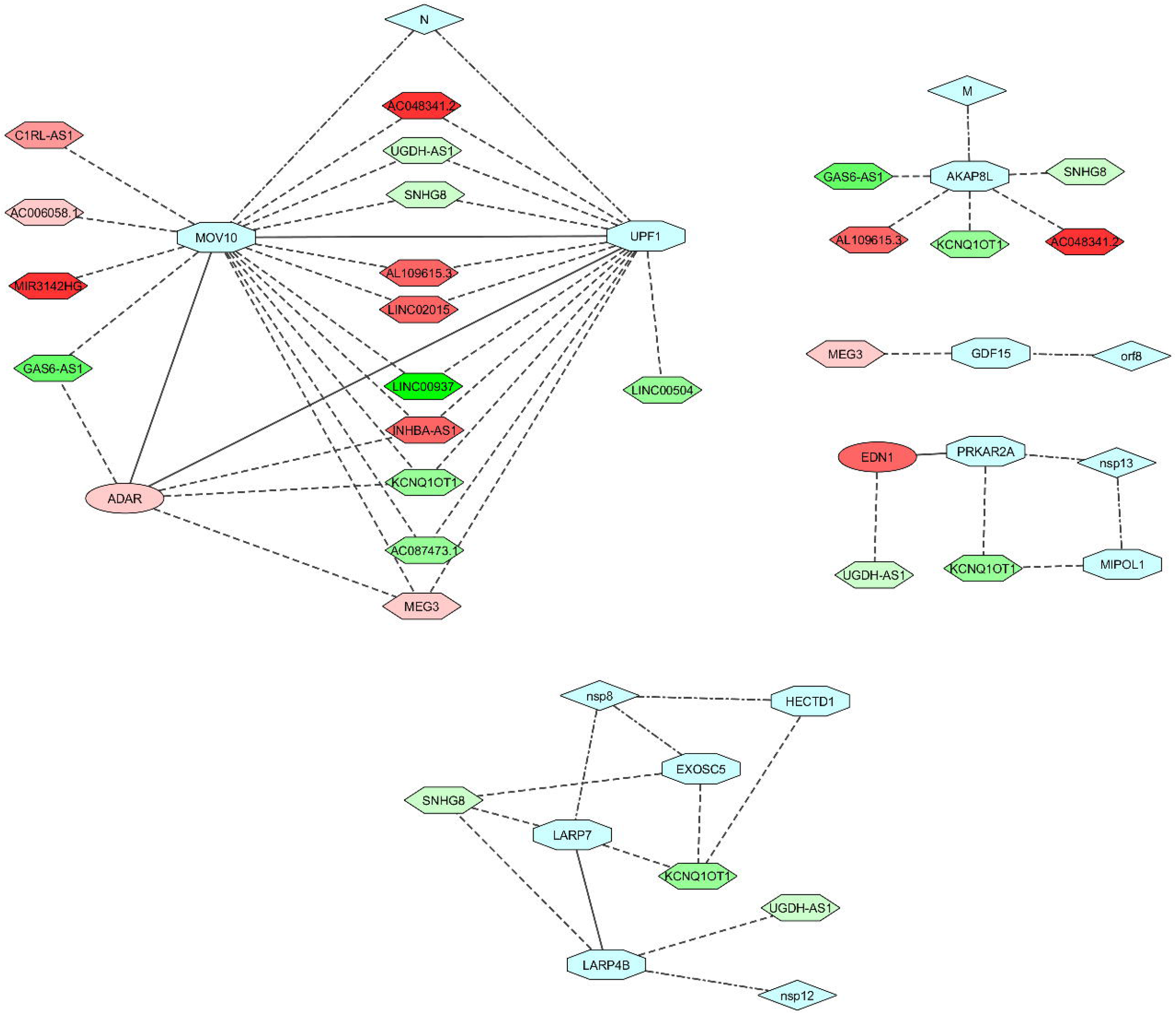
Differentially expressed lncRNAs that interact with SARS-CoV-2 protein interacting genes. Color codes, node and edge notations are similar as Figure 2.

AKAP8L, an M protein interactor, protein interacts directly with downregulated *KCNQ1OT1*, *GAS6-AS1*, *SNHG8*, *AC048341.2* and upregulated *AL109615.3* (Figure 3). Among N protein interactors, UPF1 and MOV10 proteins interacted directly with DE lncRNAs, and interacted with each other (Figure 3). UPF1 protein interacts with downregulated *AC048341.2*, *LINC00937*, *GAS6-AS1*, *KCNQ1OT1*, *UGDH-AS1*, *SNHG8*, and *AC087473.1* and upregulated *AL109615.3*, *INHBA-AS1*, *MEG3*, *LINC02015*, *MIR3142HG*, *AC006058.1* and *C1RL-AS1* (Figure 3). MOV10 protein interacts with *LINC00504*, *AC048341.2*, *LINC00937*, *GAS6-AS1*, *KCNQ1OT1*, *UGDH-AS1*, *SNHG8* and *AC087473.1*, which were downregulated, and *AL109615.3*, *INHBA-AS1*, *MEG3*, and *LINC02015*, which were upregulated (Figure 3). Both UPF1 and MOV10 proteins also interact with ADAR protein which can interact with *KCNQ1OT1*, *GAS6-AS1*, *MEG3*, and *INHBA-AS1* (Figure 3).*MEG3* can stimulate expression of *GDF15* by enhancing p53 binding to the *GDF15* gene promoter, while GDF15 can be targeted by viral orf8 (Figure 3).

Among Nsp8 interactors, EXOSC5 and LARP7 proteins interact with *KCNQ1OT1*, and *SNHG8*, both downregulated (Figure 3); *HECTD1* mRNA interacts directly with *KCNQ1OT1* (Figure 3). LARP7 also interacts with LARP4B protein, which interacts with viral Nsp12 and host lncRNA *SNHG8*, and *UGDH-AS1* (Figure 3). Among Nsp12 protein interactors, LARP4B protein interacts with *UGDH-AS1* and *SNHG8*, both downregulated (Figure 3). It also interacts with LARP7, which interacts with viral Nsp8 and host *SNHG8*, and *KCNQ1OT1* (Figure 3). Among Nsp13 interactors, *MIPOL1* and *PRKAR2A* both have RNA-RNA interaction with *KCNQ1OT1*, which is downregulated. PRKAR2A also interacts with EDN1, which interacts with *UGDH-AS1* (Figure 3).

### Upregulated targets of virally-induced microRNAs might be protected by competing lncRNAs

Competing endogenous RNA (ceRNA) hypothesis illustrates that lncRNAs and other RNA molecules harboring microRNA (miRNA) response elements can suppress the expression and biological function of each other by competing for miRNAs that can bind to the complementary regions, thus regulating miRNA-mediated gene silencing of the target genes (17, 26). Growing evidence indicates that the ceRNA regulation mechanism plays a role in disease progression and drug efficiency (27–30).

To reveal if such networks exists in SARS-CoV-2 infected cells, a list of miRNAs induced in viral infection was curated from miRwayDB database (31), Girardi et al. (32), and Leon-Icaza et al. (33). Among them, few were found to target upregulated genes, which were apparently not targeted, as DE lncRNAs were targeted simultaneously (Figure 4). The names and functions of all target genes of the selected miRNAs are provided in supplementary file 3.

**Figure 4:**
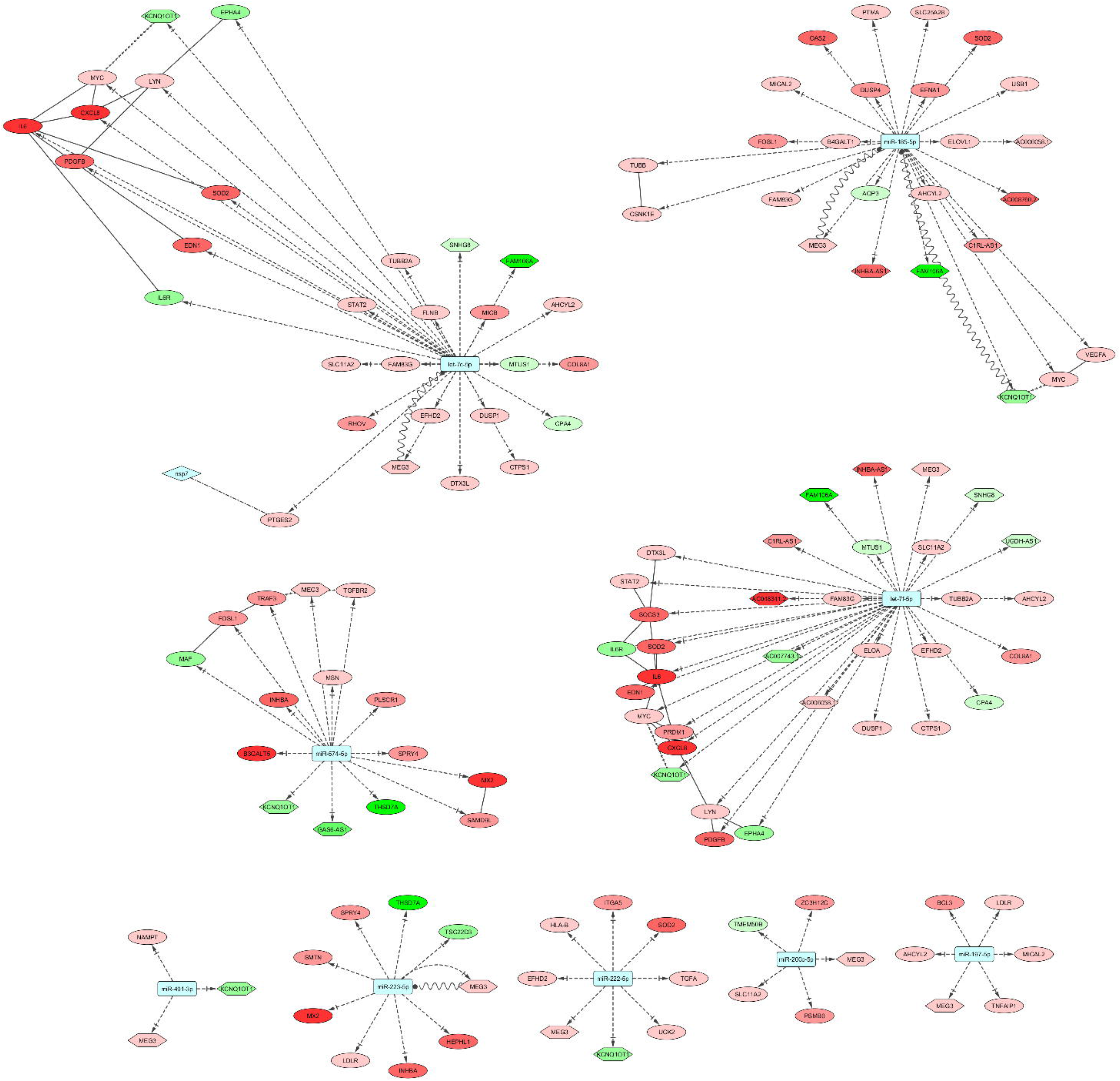
Potential virally induced microRNAs and their targets. Color depth of the node conveys level of expression. Color codes, node and edge notations are similar as Figure 2.

In SARS-CoV-2 infection, miRNA let-7c might target 21 upregulated gene and can also target *MEG3*, *KCNQ1OT1*, *SNHG8* and *FAM106A* lncRNAs (Figure 4).The 21 upregulated targets of let-7c-5p are significantly enriched for KEGG pathways like- Kaposi’s sarcoma-associated herpesvirus infection, and JAK-STAT signaling pathway (data not shown). Among the target genes, PTGES2 protein was found to interact with SARS-CoV-2 Nsp7 protein (Figure 4).

miRNA let-7f can target 20 of the upregulated genes in SARS-CoV-2 infection, including *SOCS3*. It might also target lncRNAs *AC006058.1*, *AC007743.1*, *AC048341.2*, *C1RL-AS1*, *FAM106A*, *INHBA-AS1*, *KCNQ1OT1*, *MEG3* and *SNHG8* (Figure 4). The 20 upregulated targets of the let-7f-5p are involved in pathways like- Kaposi’s sarcoma-associated herpesvirus infection related, JAK-STAT signaling and Hepatitis B related signaling (data not shown).

miR-185-5p targets 17 upregulated genes, but also targets lncRNAs *AC006058.1*, *AC008760.2*, *C1RL-AS1*, *FAM106A*, *INHBA-AS1*, *KCNQ1OT1* and *MEG3* (Figure 4). Among the 17 upregulated targets of the miR-185-5p miRNA, most significantly enriched KEGG pathways were MAPK signaling pathway and Wnt signaling pathway (data not shown). miR-197-5p, miR-200c-5p, miR-222 and miR-223-5p target 5, 3, 6 and 6 upregulated genes, respectively. They all can target lncRNA *MEG3* whilemiR-222 can additionally target *KCNQ1OT1* (Figure 4). The 6 upregulated targets of the miR-185-5p are involved in phagosome, and PI3K-Akt signaling pathway (data not shown). miR-491-3p targets upregulated *NAMPT* gene, but also targets *MEG3* lncRNA (Figure 4). miR-574-5p targets 10 upregulated genes, but also targets lncRNAs *MEG3*, *KCNQ1OT1* and *GAS6-AS1* (Figure 4). These 10 upregulated targets of the miR-574-5p are found significantly enriched in biological process/pathways like-TGF-beta signaling pathway, IL-17 signaling pathway, and apoptotic process (data not shown).

### Involvement of lncRNA may modulate important cellular pathways

Cell survival can be beneficial for continued viral replication and growth, but apoptosis is also necessary for spread. Pathways related to cell survival are targeted by virally-induced microRNAs. IL-6/JAK/STAT3 signalling acts in favor of cell survival(34). Upregulation of IL-6 during certain viral infections may promote virus survival and/or exacerbation of clinical disease (35). Among let-7c-5p targets and let-7f-5p targets, four upregulated genes, *IL6*, *MYC*, *PDGFB* and *STAT2* belong to JAK-STAT signaling pathway. *KCNQ1OT1* can act as a sponge for *MYC* gene, a target of miR-185-5p along with these miRNAs (36) (Figure 4). Also, both let-7c and let-7f target *IL6*, and absence of this interaction may lead to cell transformation progressing from initial inflammation, implying cell survival (37).

Wnt signaling pathway is known to regulate apoptosis through a variety of mechanisms (38) and it can respond to viral infections through modulation of β-catenin stabilization(39). *MYC*, *CSNK1E*, and *FOSL1* belong to Wnt signaling pathway and are targeted by miR-185-5p. *FOSL1* is also targeted by miR-574-5p (Figure 4).

Activation of PI3K-Akt signaling is a known viral strategy to delay apoptosis and prolong viral replication, as seen in acute and persistent infection (40). Three upregulated genes, *EFNA1*, *MYC* and *VEGFA*, targeted by miR-185-5p, and two upregulated genes targeted by miR-222-5p, *ITGA5* and *TGFA*, belong to PI3K-Akt signaling pathway (Figure 4). Also, miR-185 was found to inhibit cell proliferation and induce apoptosis by targeting *VEGFA* directly in von Hippel-Lindau–inactivated clear cell renal cell carcinoma (41).

Cellular response to viral infection can make or break the progression of viral life cycle. Virally-induced microRNAs are also involved in targeting pathways related with inflammation and host defense. NF-κB signaling pathway is involved in upregulating various pro-inflammatory genes that encode cytokines and chemokines (42). It is a major regulator of antiviral response and NF-κB activation pathways are manipulated by viruses to avoid cellular mechanisms that eliminate the infection (43, 44). Two upregulated gene targets of let-7c-5p and let-7f-5p, *CXCL8* and *LYN*, and two upregulated targets of miR-574-5p, *TRAF3* and *TGFBR2* is involved in NF-KappaB signaling pathway (Figure 4). *MEG3* can act as a sponge to promote upregulation of *TRAF3* (45) and *TGFBR2* (46). *BCL3*, a target of miR-197-5p, is involved in regulation of cell proliferation and participates in NF-KappaB signaling pathway (Figure 4).

MAPK pathway positively regulates virus replication in diverse group of viruses (47). 3 upregulated gene targets of let-7c-5p, *MYC*, *PDGFB*, and *DUSP1*, belong to MAPK signaling pathway. *LDLR*, involved with cholesterol homeostasis, is targeted by miR-197-5p and miR-223-5p (Figure 4).

## Discussion

Long noncoding RNAs are regulators and modulators of complex cellular pathways and viral infection is no exception (19, 20). Dysregulation of lncRNA expression and interaction, either directly with viral RNA, or indirectly with host RNA, affect the progression of viral infection (12). LncRNAs can work as a sponge for miRNA, bind protein as a competitive inhibitor, inhibit protein-protein interaction, influence post-translational modification, or affect the activity of target protein. They can modulate viral life cycle, regulate innate immune response, or assist adaptive immunity (18, 48). In our exploration of the deregulated lncRNA in SARS-CoV-2 infected NHBE cells, all of these functionalities come to fore. In our study, lncRNA-interacting DE protein-coding genes and SARS-CoV-2 protein interactors were involved in various pathways. Both RNA-RNA and RNA-protein interactions were present, indicating a complex interplay between the components involved. We also looked for possible lncRNAs exerting effects as competing endogenous RNAs to ensure a particular miRNA does not downregulate a gene. In turn, the upregulated genes play a crucial role in viral replication, disease progression, and immune response. Although some of these miRNAs also target a few downregulated genes, their decrease in expression may be controlled by separate mechanisms. Our findings shed light on the possible role of DE lncRNAs in specific processes involved with viral infection, proliferation and cellular response.

SARS-CoV-2 Nsp7-Nsp8-Nsp12 proteins interact to form a multi-subunit RNA-synthesis complex, where Nsp12 works as the RNA-dependent RNA polymerase. EXOSC5, interactor of Nsp8 protein, is a non-catalytic component of the RNA exosome complex. RNA exosome complex is involved in 3′ processing of various stable RNA species and is crucial for RNA quality control in the nucleus (49), thus Nsp8 binding may be fundamental to diminishing the capacity of exosome to act against viral mRNAs. Antiviral drug Remdesivir has been predicted to target and disassemble this complex (50, 51). Interaction of EXOSC5 protein with *KCNQ1OT1* and *SNHG8* can modulate this interaction. The putative RNA-RNA interaction between upregulated *AC006058.1* and *MALL* mRNA, a gene involved in cholesterol homeostasis and membrane trafficking, can exert an influence in maturation of SARS-CoV-2, an enveloped virus. *FAM230J*, which has no reported function, is downregulated in the infected cells. Its absence may activate the upregulated *KYNU*, which is involved in metabolite biosynthesis pathways, through lack of RNA-RNA interaction.

Host RNA-binding proteins that interact with SARS-CoV-2 proteins are involved in regulating viral transcription and mRNA stability. SARS-CoV-2 N protein interacts with UPF1 and MOV10. In case for murine hepatitis virus, a model coronavirus, the N protein carries out this interaction to inhibit nonsense-mediated decay of viral mRNAs containing multiple stop codons, thus favoring viral mRNA transcription. NMD pathways recognized cytoplasmic viral mRNAs as a substrate for degradation, but viral replication induced the inhibition of the NMD pathway through N protein (52). In human T-lymphotropic virus type 1 (HTLV-1)-infected cells, viral protein Tax bound to components of NMD pathways, including UPF1, to inhibit the process (53). SARS-CoV-2 M protein interacts with AKAP8L, which assists in viral infection progression through favoring transcription. Both LARP7 (nsp8 interactor) and LARP4B (nsp12 interactor) are RNA-binding proteins involved in RNA transcription regulation. Interaction of these RNA-binding proteins with the DE lncRNAs may be crucial for viral mRNA transcription and stability against cellular defense.

*ADAR* can be regarded as the “Editor-in-Chief” of innate immunity against viral infection (54), thus the protein’s interaction with both upregulated and downregulated lncRNAs may hold crucial role in SARS-CoV-2 replication cycle. *MEG3* was found to act as a biomarker and regulate cell functions by targeting *ADAR* in colorectal cancer (CRC). The cells overexpressing *MEG3* exhibited increased *ADAR* expression, and downregulation of *MEG3* was found in CRC tissues, cell lines and serum (55). In the infected cells, *MEG3* upregulation may lead to *ADAR* overexpression, which could have been favorable to the virus, as evidenced in Influenza A (56). *INHBA-AS1* upregulation in virus-infected cells may also contribute to cell survival through interaction with ADAR, as evidenced in gastric cancer (57) and oral squamous cell carcinoma (58).

Cell survival can be both beneficial to virus and be inhibitory. Viruses need the cell to survive certain period to undergo replication, but they also need apoptosis to exit the cell and infect others. Thus, cell survival and apoptosis becomes a key indicator of pathogenesis. Among lncRNAs, downregulated *SNHG8* has possible link to apoptosis and cell death (59), as does *KCNQ1OT1* (60–63). Unlike both, downregulation of *GAS6-AS1* may be linked with cell proliferation and survival (64, 65). All three interact with YWHAG, which is involved in signal transduction. IL-6/JAK/STAT3 signaling, Wnt signaling pathway, and PI3K-Akt signaling, involved in cell survival, are also protected by probable ceRNA function by lncRNA.

Innate immune response against the infection is inevitable, but the complex interactions underlying its activation and effect may have been the target of lncRNAs.HECTD1 is an E3 ubiquitin-protein ligase that interacts with Nsp8. As it is involved in class I MHC mediated antigen processing and presentation and innate immune system, the binding may lead to modulation of that response. In case of HECTD1 mRNA, absence of RNA-RNA interaction with downregulated *KCNQ1OT1* lncRNA can facilitate the response. PTGES2 gene was upregulated and also found to interact with SARS-CoV-2 Nsp7 protein. As it is involved in innate immune system and signaling pathways, this binding may exert an indirect influence. TLR2, as part of the innate immune response, has been connected to antiviral action against multiple viral infections (66–69). *KCNQ1OT1*, the lncRNA having putative RNA-RNA interation with TLR2, was found to attenuate sepsis-induced myocardial injury via regulating miR-192-5p/*XIAP* axis. Downregulation of the lncRNA advanced cardiac injury by allowing miR-192-5p to target *XIAP*(70). *XIAP* functions in the inhibition of apoptosis, whereas *TLR2* is known to promote apoptosis. The downregulation of *KCNQ1OT1* in infected cells thus will promote apoptosis, in accordance with its interaction with overexpressed *TLR2* mRNA. Response by NF-κB signaling pathway against the viral infection is probably protected by lncRNA against inhibition by virally-induced miRNAs.

DE lncRNAs also had probable involvement in regulating cellular processes, including signaling. The downregulation of *UGDH-AS1* can be related to *EDN1* overexpression, which can be crucial in vasoconstriction, leading to severe symptoms, as seen in SARS-CoV-2 infections (71). SARS-CoV-2 Nsp13 protein works as a viral helicase, and interacts with MIPOL1 and PRKAR2A. Function of MIPOL1 is unclear, whereas PRKAR2A is involved in cAMP-mediated signaling, which is the probable target for modulation by Nsp13. The RNA-RNA interactions of downregulated *KCNQ1OT1* with the mRNAs of these two genes may have implications on viral infection progression. Additionally, MAPK pathway is probably regulated through lncRNA involvement.

From literature, it was confirmed that *KCNQ1OT1* (72) and *MEG3* (73) both can act as probable ceRNA targets for miR-185-5p, whereas *MEG3* can act as a ceRNA for let-7c-5p (74). This proves the existence of an expanded role of these lncRNAs in infected cells.

## Conclusions

Our findings explore the molecular footprint of SARS-CoV-2 infection from the lens of lncRNAs. The integrated analysis has linked multiple actors in the complex interplay of molecules in exacerbating the infection and identified possible drug targets. We trace the involvement of lncRNAs with cellular behavior in this situation and illuminate their role in cell survival, viral replicaion, and immune defense. The lethality and swift transmission of this virus is entwined with the deregulated cellular environment, and lncRNA regulation is crucial for understanding its parameters. Our results could provide insights for scheming some novel RNA therapeutics during the lacking of effective cure option.

## Methods

### Identification of differentially expressed lncRNAs

Illumina sequenced RNA-seq raw FastQ reads were extracted from GEO database accession: GSE147507 (75). This data include independent biological triplicates of primary human lung epithelium (NHBE) which were mock treated or infected with SARS-CoV-2 for 24hrs. Mapping of reads was done with TopHat (tophat v2.1.1 with Bowtie v2.4.1) (76). Short reads were uniquely aligned allowing at best two mismatches to the latest version human reference genome from (GRCh38) as downloaded from USCS database (77). Sequence matched exactly more than one place with equally quality were discarded to avoid bias (78). The reads that were not mapped to the genome were utilized to map against the transcriptome (junctions mapping). Latest version of Ensembl gene model (79) (version 99, as extracted from UCSC) was used for this process. After mapping, we used SubRead package featureCount v2.21 (80) to calculate absolute read abundance (read count, rc) for each transcript/gene associated to the Ensembl genes. For differential expression (DE) analysis we used DESeq2 v1.26.0 with R v3.6.2 (2019-07-05) (81) that uses a model based on the negative binomial distribution. To avoid false positive, we considered only those transcripts where at least 10 reads were annotated in at least one of the samples used in this study.

### Protein-protein interaction (PPI) network construction

From RNA-seq data analysis, identified differentially expressed 638 protein-coding genes and recently reported 332 SARS-CoV-2 protein interactors by Gordon et al. (25) were used in combination to build PPI network. Edge information for the network built with these proteins were extracted from the STRING (82) database. Interactions with high confidence (score>0.700) were selected and retrieved. A network file was prepared in SIF format to be visualized using Cytoscape v3.7.2 (83).

### Retrieval of RNA-RNA interactions and lncRNA functions

RNA-RNA interactions between the DE lncRNAs and other RNAs were retrieved from NPInter v4.0 (84). The interacting RNAs were searched for identified DE protein-coding genes and viral protein-interactor genes. Functions of the DE lncRNAs were retrieved from LncBook (85). The viral proteins that bind to proteins with lncRNA interactors were also identified.

### Identification of virally-induced miRNAs

miRwayDB(31) was utilized to look for miRNAs involved in viral-mediated disease. Additionally, host miRNAs involved in viral infection as identified by Girardi et al. (32) and microRNAs in viral acute respiratory infections as described by Leon-Icaza et al. (33) were retrieved. These miRNAs were manually curated for their involvement in viral pathogenesis and possible connection to SARS-CoV-2.

### Extraction of microRNA targets

Gene targets for the curated set of miRNAs were retrieved from experimentally validated miRTarBase (86) and filtered for DE protein-coding genes. DIANA-LncBase v3 (87) was used to retrieve miRNAs that target the DE lncRNAs, with high-confidence interactions considered only. The lists were combined to identify the miRNAs that target both DE protein-coding genes and lncRNAs. The set of upregulated target genes for each miRNA were analyzed using NetworkAnalyst 3.0 (88) tools for functional overrepresentation and network enrichment. miRNAs that mostly targeted upregulated genes were finally selected.

### Construction of biological networks

Networks denoting the interactions between DE lncRNAs, DE genes, viral proteins, and curated miRNAs were built using Cytoscape v3.7.2 (83).

## Declarations

### Ethics approval and consent to participate

Not applicable.

### Consent for publication

Not applicable.

### Availability of data and materials

Publicly available data were utilized. Analyses generated data are deposited as supplementary files.

### Authors’ Contributions

ABMMKI conceived the project. ABMMKI and RRT designed the workflow. RRT, MAAKK and ABMMKI performed the analyses. RRT, MAAKK and ABMMKI wrote the manuscript. All authors read and approved the final manuscript.

## Acknowledgements

Not applicable.

## List of supplementary files

**Supplementary file 1**: List of differentially expressed genes in SARS-CoV-2 infected cells.

**Supplementary file 2**: List of Human proteins and their associated interacting SARS-CoV-2 proteins.

**Supplementary file 3**: DE Genes targeted by virally induced microRNAs.

