## Supplementary file 3 for "Perversely expressed long noncoding RNAs can alter host response and viral proliferation in SARS-CoV-2 infection"

**Supplementary file 3: DE Genes targeted by virally induced microRNAs.**

| **Gene Name** | **Protein Function** | **Targeting MicroRNA** |
| --- | --- | --- |
| *AHCYL2* | Adenosylhomocysteinase 3; May regulate the electrogenic sodium/bicarbonate cotransporter SLC4A4 activity and Mg(2+)-sensitivity. | miR-185-5p  miR-197-5p  let-7c-5p  let-7f-5p |
| *AQP3* | Aquaporin-3; Water channel required to promote glycerol permeability and water transport across cell membranes. Acts as a glycerol transporter in skin and plays an important role in regulating SC (stratum corneum) and epidermal glycerol content. Involved in skin hydration, wound healing, and tumorigenesis. | miR-185-5p |
| *B3GALT5* | Beta-1,3-galactosyltransferase 5; Catalyzes the transfer of Gal to GlcNAc-based acceptors with a preference for the core3 O-linked glycan GlcNAc(beta1,3)GalNAc structure. | miR-574-5p |
| *B4GALT1* | Beta-1,4-galactosyltransferase 1; The Golgi complex form catalyzes the production of lactose in the lactating mammary gland and could also be responsible for the synthesis of complex-type N-linked oligosaccharides in many glycoproteins. | miR-185-5p |
| *BCL3* | B-cell lymphoma 3 protein; Contributes to the regulation of transcriptional activation of NF-kappa-B target genes. In the cytoplasm, inhibits the nuclear translocation of the NF-kappa-B p50 subunit. In the nucleus, acts as transcriptional activator that promotes transcription of NF-kappa-B target genes. | miR-197-5p |
| *COL8A1* | Collagen alpha-1(VIII) chain; Macromolecular component of the subendothelium. Major component of the Descemet's membrane (basement membrane) of corneal endothelial cells. Also component of the endothelia of blood vessels. | let-7c-5p  let-7f-5p |
| *CPA4* | Carboxypeptidase A4; Metalloprotease that could be involved in the histone hyperacetylation pathway. Releases a C-terminal amino acid, with preference for -Phe, -Leu, -Ile, -Met, -Tyr and –Val. | let-7c-5p  let-7f-5p |
| *CSNK1E* | Casein kinase I isoform epsilon; Casein kinases are operationally defined by their preferential utilization of acidic proteins such as caseins as substrates. Can phosphorylate a large number of proteins. Participates in Wnt signaling. Phosphorylates DVL1 and DVL2. Central component of the circadian clock. | miR-185-5p |
| *CTPS1* | CTP synthase 1; This enzyme is involved in the de novo synthesis of CTP, a precursor of DNA, RNA and phospholipids. Catalyzes the ATP- dependent amination of UTP to CTP with either L-glutamine or ammonia as a source of nitrogen. | let-7c-5p  let-7f-5p |
| *CXCL8* | Interleukin-8; IL-8 is a chemotactic factor that attracts neutrophils, basophils, and T-cells, but not monocytes. It is also involved in neutrophil activation. It is released from several cell types in response to an inflammatory stimulus. | let-7c-5p  let-7f-5p |
| *DTX3L* | E3 ubiquitin-protein ligase DTX3L; E3 ubiquitin-protein ligase which, in association with ADP-ribosyltransferase PARP9, plays a role in DNA damage repair and in interferon-mediated antiviral responses. Monoubiquitinates several histones, including histone H2A, H2B, H3 and H4. In response to DNA damage, mediates monoubiquitination of 'Lys-91' of histone H4 (H4K91ub1). | let-7c-5p  let-7f-5p |
| *DUSP1* | Dual specificity protein phosphatase 1; Dual specificity phosphatase that dephosphorylates MAP kinase MAPK1/ERK2 on both 'Thr-183' and 'Tyr-185', regulating its activity during the meiotic cell cycle. | let-7c-5p  let-7f-5p |
| *DUSP4* | Dual specificity protein phosphatase 4; Regulates mitogenic signal transduction by dephosphorylating both Thr and Tyr residues on MAP kinases ERK1 and ERK2 | miR-185-5p |
| *EDN1* | Endothelin-1; Endothelins are endothelium-derived vasoconstrictor peptides; Belongs to the endothelin/sarafotoxin family | let-7c-5p  let-7f-5p |
| *EFHD2* | EF-hand domain-containing protein D2; May regulate B-cell receptor (BCR)-induced immature and primary B-cell apoptosis. Plays a role as negative regulator of the canonical NF-kappa-B-activating branch. Controls spontaneous apoptosis through the regulation of BCL2L1 abundance. | miR-222-5p  let-7c-5p  let-7f-5p |
| *EFNA1* | Ephrin-A1; Cell surface GPI-bound ligand for Eph receptors, a family of receptor tyrosine kinases which are crucial for migration, repulsion and adhesion during neuronal, vascular and epithelial development. Binds promiscuously Eph receptors residing on adjacent cells, leading to contact-dependent bidirectional signaling into neighboring cells. Plays an important role in angiogenesis and tumor neovascularization. | miR-185-5p |
| *ELOVL1* | Elongation of very long chain fatty acids protein 1; Catalyzes the first and rate-limiting reaction of the four that constitute the long-chain fatty acids elongation cycle. This endoplasmic reticulum-bound enzymatic process, allows the addition of 2 carbons to the chain of long- and very long-chain fatty acids/VLCFAs per cycle. | miR-185-5p |
| *EPHA4* | Ephrin type-A receptor 4; Receptor tyrosine kinase which binds membrane-bound ephrin family ligands residing on adjacent cells, leading to contact-dependent bidirectional signaling into neighboring cells. Highly promiscuous, it has the unique property among Eph receptors to bind and to be physiologically activated by both GPI-anchored ephrin-A and transmembrane ephrin-B ligands including EFNA1 and EFNB3. | let-7c-5p  let-7f-5p |
| *FAM83G* | Protein FAM83G; May regulate the bone morphogenetic proteins (BMP) pathway | miR-185-5p  let-7c-5p  let-7f-5p |
| *FLNB* | Filamin-B; Connects cell membrane constituents to the actin cytoskeleton. May promote orthogonal branching of actin filaments and links actin filaments to membrane glycoproteins. Anchors various transmembrane proteins to the actin cytoskeleton. Interaction with FLNA may allow neuroblast migration from the ventricular zone into the cortical plate. | let-7c-5p |
| *FOSL1* | Fos-related antigen 1; FOS like 1, AP-1 transcription factor subunit; Belongs to the bZIP family. Fos subfamily | miR-185-5p  miR-574-5p |
| *HEPHL1* | Hephaestin-like protein 1; May function as a ferroxidase and may be involved in copper transport and homeostasis | miR-223-5p |
| *HLA-B* | HLA class I histocompatibility antigen, B-7 alpha chain; Involved in the presentation of foreign antigens to the immune system; C1-set domain containing | miR-222-5p |
| *IL6* | Interleukin-6; Cytokine with a wide variety of biological functions. It is a potent inducer of the acute phase response. Plays an essential role in the final differentiation of B-cells into Ig- secreting cells Involved in lymphocyte and monocyte differentiation. Acts on B-cells, T-cells, hepatocytes, hematopoietic progenitor cells and cells of the CNS. Required for the generation of T(H)17 cells. Also acts as a myokine. | let-7c-5p  let-7f-5p |
| *IL6R* | Interleukin-6 receptor subunit alpha; Part of the receptor for interleukin 6. Binds to IL6 with low affinity, but does not transduce a signal. Signal activation necessitate an association with IL6ST. Activation may lead to the regulation of the immune response, acute-phase reactions and hematopoiesis; CD molecules | let-7c-5p  let-7f-5p |
| *INHBA* | Inhibin beta A chain; Inhibins and activins inhibit and activate, respectively, the secretion of follitropin by the pituitary gland. Inhibins/activins are involved in regulating a number of diverse functions such as hypothalamic and pituitary hormone secretion, gonadal hormone secretion, germ cell development and maturation, erythroid differentiation, insulin secretion, nerve cell survival, embryonic axial development or bone growth, depending on their subunit composition. | miR-574-5p  miR-223-5p |
| *ITGA5* | Integrin alpha-5; Integrin alpha-5/beta-1 is a receptor for fibronectin and fibrinogen. It recognizes the sequence R-G-D in its ligands. ITGA5:ITGB1 binds to PLA2G2A via a site (site 2) which is distinct from the classical ligand-binding site (site 1) and this induces integrin conformational changes and enhanced ligand binding to site 1. ITGA5:ITGB1 acts as a receptor for fibrillin-1 (FBN1) and mediates R-G-D-dependent cell adhesion to FBN1; CD molecules | miR-222-5p |
| *LDLR* | Low-density lipoprotein receptor; Binds LDL, the major cholesterol-carrying lipoprotein of plasma, and transports it into cells by endocytosis. In order to be internalized, the receptor-ligand complexes must first cluster into clathrin-coated pits; Belongs to the LDLR family | miR-197-5p  miR-223-5p |
| *LYN* | Tyrosine-protein kinase Lyn; Non-receptor tyrosine-protein kinase that transmits signals from cell surface receptors and plays an important role in the regulation of innate and adaptive immune responses, hematopoiesis, responses to growth factors and cytokines, integrin signaling, but also responses to DNA damage and genotoxic agents. | let-7c-5p  let-7f-5p |
| *MAF* | Transcription factor Maf; Acts as a transcriptional activator or repressor. Involved in embryonic lens fiber cell development. Recruits the transcriptional coactivators CREBBP and/or EP300 to crystallin promoters leading to up-regulation of crystallin gene during lens fiber cell differentiation. Activates the expression of IL4 in T helper 2 (Th2) cells. Increases T-cell susceptibility to apoptosis by interacting with MYB and decreasing BCL2 expression. | miR-574-5p |
| *MICAL2* | [F-actin]-monooxygenase MICAL2; Nuclear monooxygenase that promotes depolymerization of F-actin by mediating oxidation of specific methionine residues on actin to form methionine-sulfoxide, resulting in actin filament disassembly and preventing repolymerization. In the absence of actin, it also functions as a NADPH oxidase producing H(2)O(2) (By similarity). Acts as a key regulator of the SRF signaling pathway elicited by nerve growth factor and serum. | miR-185-5p  miR-197-5p |
| *MICB* | MHC class I polypeptide-related sequence B; Seems to have no role in antigen presentation. Acts as a stress-induced self-antigen that is recognized by gamma delta T cells. Ligand for the KLRK1/NKG2D receptor. Binding to KLRK1 leads to cell lysis. | let-7c-5p |
| *MSN* | Moesin; Probably involved in connections of major cytoskeletal structures to the plasma membrane. May inhibit herpes simplex virus 1 infection at an early stage. Plays a role in regulating the proliferation, migration, and adhesion of human lymphoid cells and participates in immunologic synapse formation; FERM domain containing | miR-574-5p |
| *MTUS1* | Microtubule-associated tumor suppressor 1; Cooperates with AGTR2 to inhibit ERK2 activation and cell proliferation. May be required for AGTR2 cell surface expression. Together with PTPN6, induces UBE2V2 expression upon angiotensin-II stimulation. | let-7c-5p  let-7f-5p |
| *MX2* | Interferon-induced GTP-binding protein Mx2; Interferon-induced dynamin-like GTPase with potent antiviral activity against human immunodeficiency virus type 1 (HIV-1). Acts by targeting the viral capsid and affects the nuclear uptake and/or stability of the HIV-1 replication complex and the subsequent chromosomal integration of the proviral DNA. Exhibits antiviral activity also against simian immunodeficiency virus (SIV-mnd). | miR-574-5p  miR-223-5p |
| *MYC* | Myc proto-oncogene protein; Transcription factor that binds DNA in a non-specific manner, yet also specifically recognizes the core sequence 5'- CAC[GA]TG-3'. Activates the transcription of growth-related genes. Binds to the VEGFA promoter, promoting VEGFA production and subsequent sprouting angiogenesis. | miR-185-5p  let-7c-5p  let-7f-5p |
| *NAMPT* | Nicotinamide phosphoribosyltransferase; Catalyzes the condensation of nicotinamide with 5- phosphoribosyl-1-pyrophosphate to yield nicotinamide mononucleotide, an intermediate in the biosynthesis of NAD. It is the rate limiting component in the mammalian NAD biosynthesis pathway. The secreted form behaves both as a cytokine with immunomodulating properties and an adipokine with anti-diabetic properties. | miR-491-3p |
| *OAS2* | 2'-5'-oligoadenylate synthase 2; Interferon-induced, dsRNA-activated antiviral enzyme which plays a critical role in cellular innate antiviral response. In addition, it may also play a role in other cellular processes such as apoptosis, cell growth, differentiation and gene regulation. Synthesizes higher oligomers of 2'-5'-oligoadenylates (2-5A) from ATP which then bind to the inactive monomeric form of ribonuclease L (RNase L) leading to its dimerization and subsequent activation. Activation of RNase L leads to degradation of cellular as well as viral RNA, resulting in the inhibition of viral infection. | miR-185-5p |
| *PDGFB* | Platelet-derived growth factor subunit B; Growth factor that plays an essential role in the regulation of embryonic development, cell proliferation, cell migration, survival and chemotaxis. Potent mitogen for cells of mesenchymal origin. Required for normal proliferation and recruitment of pericytes and vascular smooth muscle cells in the central nervous system, skin, lung, heart and placenta. Plays an important role in wound healing. | let-7c-5p  let-7f-5p |
| *PLSCR1* | Phospholipid scramblase 1; May mediate accelerated ATP-independent bidirectional transbilayer migration of phospholipids upon binding calcium ions that results in a loss of phospholipid asymmetry in the plasma membrane. May play a central role in the initiation of fibrin clot formation, in the activation of mast cells and in the recognition of apoptotic and injured cells by the reticuloendothelial system. | miR-574-5p |
| *PRDM1* | PR domain zinc finger protein 1; Transcription factor that mediates a transcriptional program in various innate and adaptive immune tissue-resident lymphocyte T cell types such as tissue-resident memory T (Trm), natural killer (trNK) and natural killer T (NKT) cells and negatively regulates gene expression of proteins that promote the egress of tissue-resident T-cell populations from non-lymphoid organs. | let-7f-5p |
| *PSMB9* | Proteasome subunit beta type-9; The proteasome is a multicatalytic proteinase complex which is characterized by its ability to cleave peptides with Arg, Phe, Tyr, Leu, and Glu adjacent to the leaving group at neutral or slightly basic pH. The proteasome has an ATP-dependent proteolytic activity. This subunit is involved in antigen processing to generate class I binding peptides. Replacement of PSMB6 by PSMB9 increases the capacity of the immunoproteasome to cleave model peptides after hydrophobic and basic residues. | miR-200c-5p |
| *PTGES2* | Prostaglandin E synthase 2; Isomerase that catalyzes the conversion of PGH2 into the more stable prostaglandin E2 (PGE2). | let-7c-5p |
| *PTMA* | Prothymosin alpha; Prothymosin alpha may mediate immune function by conferring resistance to certain opportunistic infections. | miR-185-5p |
| *RHOV* | Rho-related GTP-binding protein RhoV; Plays a role in the control of the actin cytoskeleton via activation of the JNK pathway; Belongs to the small GTPase superfamily. | let-7c-5p |
| *SAMD9L* | Sterile alpha motif domain-containing protein 9-like; May be involved in endosome fusion. Mediates down- regulation of growth factor signaling via internalization of growth factor receptors. | miR-574-5p |
| *SLC11A2* | Natural resistance-associated macrophage protein 2; Important in metal transport, in particular iron. Can also transport manganese, cobalt, cadmium, nickel, vanadium and lead. Involved in apical iron uptake into duodenal enterocytes. Involved in iron transport from acidified endosomes into the cytoplasm of erythroid precursor cells. | miR-200c-5p  let-7c-5p  let-7f-5p |
| *SLC25A28* | Mitoferrin-2; Mitochondrial iron transporter that mediates iron uptake. Probably required for heme synthesis of hemoproteins and Fe-S cluster assembly in non-erythroid cells. The iron delivered into the mitochondria, presumably as Fe(2+), is then probably delivered to ferrochelatase to catalyze Fe(2+) incorporation into protoprophyrin IX to make heme (By similarity). | miR-185-5p |
| *SMTN* | Smoothelin; Structural protein of the cytoskeleton; Belongs to the smoothelin family | miR-223-5p |
| *SOCS3* | Suppressor of cytokine signaling 3; SOCS family proteins form part of a classical negative feedback system that regulates cytokine signal transduction. SOCS3 is involved in negative regulation of cytokines that signal through the JAK/STAT pathway. Inhibits cytokine signal transduction by binding to tyrosine kinase receptors including gp130, LIF, erythropoietin, insulin, IL12, GCSF and leptin receptors. Binding to JAK2 inhibits its kinase activity. Suppresses fetal liver erythropoiesis. | let-7f-5p |
| *SOD2* | Superoxide dismutase [14], mitochondrial; Destroys superoxide anion radicals which are normally produced within the cells and which are toxic to biological systems | miR-185-5p  miR-222-5p  let-7c-5p  let-7f-5p |
| *SPRY4* | Protein sprouty homolog 4; Suppresses the insulin receptor and EGFR-transduced MAPK signaling pathway, but does not inhibit MAPK activation by a constitutively active mutant Ras. Probably impairs the formation of GTP-Ras. | miR-574-5p  miR-223-5p |
| *STAT2* | Signal transducer and activator of transcription 2; Signal transducer and activator of transcription that mediates signaling by type I IFNs (IFN-alpha and IFN-beta). Following type I IFN binding to cell surface receptors, Jak kinases (TYK2 and JAK1) are activated, leading to tyrosine phosphorylation of STAT1 and STAT2. The phosphorylated STATs dimerize, associate with IRF9/ISGF3G to form a complex termed ISGF3 transcription factor, that enters the nucleus. ISGF3 binds to the IFN stimulated response element (ISRE) to activate the transcription of interferon stimulated genes, which drive response to viral infection. | let-7c-5p  let-7f-5p |
| *TCEB3 (ELOA)* | Elongin-A; SIII, also known as elongin, is a general transcription elongation factor that increases the RNA polymerase II transcription elongation past template-encoded arresting sites. Subunit A is transcriptionally active and its transcription activity is strongly enhanced by binding to the dimeric complex of the SIII regulatory subunits B and C (elongin BC complex). | let-7f-5p |
| *TGFA* | Protransforming growth factor alpha; TGF alpha is a mitogenic polypeptide that is able to bind to the EGF receptor/EGFR and to act synergistically with TGF beta to promote anchorage-independent cell proliferation in soft agar | miR-222-5p |
| *TGFBR2* | TGF-beta receptor type-2; Transmembrane serine/threonine kinase forming with the TGF-beta type I serine/threonine kinase receptor, TGFBR1, the non- promiscuous receptor for the TGF-beta cytokines TGFB1, TGFB2 and TGFB3. Transduces the TGFB1, TGFB2 and TGFB3 signal from the cell surface to the cytoplasm and is thus regulating a plethora of physiological and pathological processes including cell cycle arrest in epithelial and hematopoietic cells, control of mesenchymal cell proliferation and differentiation, wound healing, extracellular matrix production, immunosuppression etc. | miR-574-5p |
| *THSD7A* | Thrombospondin type-1 domain-containing protein 7A; The soluble form promotes endothelial cell migration and filopodia formationduring angiogenesis via a FAK-dependent mechanism | miR-574-5p  miR-223-5p |
| *TMEM50B* | Transmembrane protein 50B | miR-200c-5p |
| *TNFAIP1* | BTB/POZ domain-containing adapter for CUL3-mediated RhoA degradation protein 2; Substrate-specific adapter of a BCR (BTB-CUL3-RBX1) E3 ubiquitin-protein ligase complex involved in regulation of cytoskeleton structure. The BCR(TNFAIP1) E3 ubiquitin ligase complex mediates the ubiquitination of RHOA, leading to its degradation by the proteasome, thereby regulating the actin cytoskeleton and cell migration. Its interaction with RHOB may regulate apoptosis. | miR-197-5p |
| *TRAF3* | TNF receptor-associated factor 3; Regulates pathways leading to the activation of NF- kappa-B and MAP kinases, and plays a central role in the regulation of B-cell survival. Part of signaling pathways leading to the production of cytokines and interferon. Required for normal antibody isotype switching from IgM to IgG. Plays a role in T-cell dependent immune responses. Plays a role in the regulation of antiviral responses. Is an essential constituent of several E3 ubiquitin-protein ligase complexes. | miR-574-5p |
| *TSC22D3* | TSC22 domain family protein 3; Protects T-cells from IL2 deprivation-induced apoptosis through the inhibition of FOXO3A transcriptional activity that leads to the down-regulation of the pro-apoptotic factor BCL2L11. In macrophages, plays a role in the anti-inflammatory and immunosuppressive effects of glucocorticoids and IL10. In T-cells, inhibits anti-CD3-induced NFKB1 nuclear translocation. In vitro, suppresses AP1 and NFKB1 DNA-binding activities (By similarity). | miR-223-5p |
| *TUBB* | Tubulin beta chain; Tubulin is the major constituent of microtubules. It binds two moles of GTP, one at an exchangeable site on the beta chain and one at a non-exchangeable site on the alpha chain. | miR-185-5p |
| *TUBB2A* | Tubulin beta-2A chain; Tubulin is the major constituent of microtubules. It binds two moles of GTP, one at an exchangeable site on the beta chain and one at a non-exchangeable site on the alpha chain (By similarity). | let-7c-5p  let-7f-5p |
| *UCK2* | Uridine-cytidine kinase 2; Phosphorylates uridine and cytidine to uridine monophosphate and cytidine monophosphate. Does not phosphorylate deoxyribonucleosides or purine ribonucleosides. Can use ATP or GTP as a phosphate donor. Can also phosphorylate cytidine and uridine nucleoside analogs. | miR-222-5p |
| *USB1* | U6 snRNA phosphodiesterase; Phosphodiesterase responsible for the U6 snRNA 3' end processing. Acts as an exoribonuclease (RNase) responsible for trimming the poly(U) tract of the last nucleotides in the pre-U6 snRNA molecule, leading to the formation of mature U6 snRNA 3' end-terminated with a 2',3'-cyclic phosphate. | miR-185-5p |
| *VEGFA* | Vascular endothelial growth factor A; Growth factor active in angiogenesis, vasculogenesis and endothelial cell growth. Induces endothelial cell proliferation, promotes cell migration, inhibits apoptosis and induces permeabilization of blood vessels. Binds to the FLT1/VEGFR1 and KDR/VEGFR2 receptors, heparan sulfate and heparin. NRP1/Neuropilin-1 binds isoforms VEGF-165 and VEGF-145. Isoform VEGF165B binds to KDR but does not activate downstream signaling pathways, does not activate angiogenesis and inhibits tumor growth. | miR-185-5p |
| *ZC3H12C* | Probable ribonuclease ZC3H12C; May function as RNase and regulate the levels of target RNA species; Belongs to the ZC3H12 family | miR-200c-5p |
